# Dysfunctional and compensatory brain networks underlying math fluency

**DOI:** 10.1101/752089

**Authors:** Michelle AN La, Debjani Saha, Karen F Berman, Hao Yang Tan

## Abstract

Poor math fluency, or timed calculation (TC) performance, is a characteristic of dyscalculia, a common cause of poor educational and occupational outcomes. Here, we examined neural substrates of dysfunctional math fluency and potential compensatory mechanisms. We performed functional MRI scans of participants with divergent performance on an event-related TC paradigm (poor TC, <0.5 accuracy, n=34; vs. controls, accuracy>0.8, n=34). Individuals with poor TC had decreased intraparietal sulcus (IPS) engagement, and decreased IPS-striatal and IPS-prefrontal effective connectivity. We next examined an independent well-performing sample (TC accuracy>0.8, n=100), stratified according to relatively low-versus high-IPS activation during TC. Relatively reduced IPS engagement, or patterns of IPS-related effective connectivity similar to those with poor TC, appeared to be compensated for by increased engagement of effective connectivity involving fusiform gyrus, angular gyrus, inferior frontal gyrus and striatum. Neural connectivity involving high-level visual processing in fusiform gyrus and related ventral cortical networks may be relevant in compensatory function ameliorating aspects of dyscalculia and mathematical difficulty.

## Introduction

Mathematical ability plays an important part of everyday life, impacting one’s education, decision-making, employment, and wellbeing (Banks et al., 2011; Brooks and Pui, 2010; Gerardi et al., 2013; Gross, 2009; Parsons and Bynner, 2005; Reyna et al., 2009). Dyscalculia is characterized by difficulty with arithmetic and has an estimated population prevalence of 5-7% (Shalev, 2007; Shalev and Gross-Tsur, 2001). Understanding brain function associated with this manner of dysfunction, and identifying neural networks engaged in behavioral compensation may be valuable in strategies to mitigate this cognitive disability.

Multiple aspects of number processing, including arithmetic calculations, engage the intraparietal sulcus (IPS) (Ansari, 2007; Ansari et al., 2006; Cantlon et al., 2006; Cohen Kadosh et al., 2007; Dehaene et al., 2003; Eger et al., 2003; Gobell et al., 2006; Pinel et al., 2004; Venkatraman et al., 2005), and this region is implicated in numerical deficits and dyscalculia (Butterworth, 1999; Butterworth and Kovas, 2013; Landerl et al., 2004). The DSM-V categorizes dyscalculia as mathematical impairment under specific learning disorder. Mathematical learning disability (MLD) is a related classification of poor mathematical skill comprising deficits in numerical cognition, working memory, visuospatial processing, and attention (Kaufmann and von Aster, 2012; Rubinsten and Henik, 2009). Features of MLD are present in genetic disorders affecting cognition and IPS development, such as Turner syndrome, Williams syndrome, and the 22q11.2 hemi-deletion syndrome (Barnea-Goraly et al., 2005; Carvalho et al., 2014; De Smedt et al., 2009; Molko et al., 2003; O’Hearn and Luna, 2009; Quintero et al., 2014; Simon et al., 2008).

Math fluency (MF), or performance in timed calculation (TC) tasks focusing on rapid quantity and numerical manipulation, is fundamental to mathematical ability. MF is an independent building block upon which more complex mathematical problem-solving involving individualized step-by-step strategies may proceed (Petrill et al., 2012; Wang et al., 2016). MF also involves aspects of innate number sense and working memory, which contributes to variation in math ability, as distinct from untimed math performance (Fuchs et al., 2010; Galeano Weber et al., 2016; Hart et al., 2010; Mazzocco et al., 2008; Stoianov, 2014). A study of twins has suggested that untimed mathematics performance is more closely influenced by the childhood learning environment, while timed performance is more strongly influenced by genetic factors (Hart et al., 2009). MF engages number sense, or intrinsic magnitude representations mapping in the IPS (Nieder, 2016). The IPS may also limit the number of items that can be retained in working memory, affecting precision and variability of MF performance (Galeano Weber et al., 2016; McLean and Hitch, 1999).

While the IPS is fundamental to MF, less is known about the neural connectivity networks implicated in IPS engagement, or about the network functions potentially compensating for reduced IPS engagement. Functional magnetic resonance imaging (fMRI) suggests that the IPS, the angular gyrus (AG), dorsolateral prefrontal cortex (DLPFC), fusiform gyrus (FFG) and inferior frontal gyrus (IFG) are engaged in MF and calculation tasks (Arsalidou and Taylor, 2011; Dehaene et al., 2004). However, effective connectivity across these nodes and varying patterns of neural engagement cross individuals are less well characterized. Variations in IPS-related neural activation and effective connectivity patterns could inform us about potential compensatory pathways that could mitigate behavioral and neural dysfunction in IPS and MF. Thus, we sought to examine behavioral dysfunction and potential compensatory neural functions across these multiple brain regions as they relate to MF. Specifically, in experiment one, we identified brain network connectivity associated with poor MF. In experiment two, we explored an independent sample of individuals with high MF across varying activity of the IPS, with the goal of examining potential networks that may compensate for reduced engagement of IPS traditionally implicated in MF. Apart from activity at IPS, we also examined connectivity patterns compensating for IPS-related connectivity deficits associated with poor MF.

## Methods

### Participants

All participants (N=168) were healthy subjects enrolled as part of the Clinical Brain Disorders Branch Sibling Study (Egan et al., 2001). All participants were between 18 and 45 years of age and right-handed. They were interviewed by a research psychiatrist using the Structured Clinical Interview for DSM-IV to determine the presence of any psychiatric illnesses, and they completed a neurological examination and a battery of neuropsychological tests. Exclusion criteria included an IQ <70, a history of prolonged substance abuse or significant medical problems, and any abnormalities found by EEG or MRI. All participants gave written consent before participation and were reimbursed for their time. The study was approved by the NIMH Institutional Review Board.

### Cognitive paradigm

Blood oxygen level-dependent functional MRI data were acquired from subjects as they performed number-related tasks after a brief training period (∼10 min). The event-related paradigm was described in detail in a previous study (Tan et al., 2007). Briefly, subjects performed simple arithmetic and number size comparisons on two single digit integers during a session of 8 min (Figure 1). Specifically, they performed an arithmetic subtraction of 2 or 3 from one of two integers presented, and were subsequently prompted to identify with a left or right button press the larger result, or in an equal number of randomly spaced trials, the smaller result. These timed calculation task events (16 total) included performing arithmetic (computation and size judgment, CJ, 8 trials), or doing this after first encoding two digits in working memory (encoding, followed by the same computation, and size judgment, E_CJ, 8 trials, Figure 1). An equal number of control tasks which did not engage arithmetic were also administered, i.e. comparing the relative size of two integers with a similar working memory maintenance load after encoding (16 trials), and a simple motor task where subjects followed instructions to press the left or right button. A jittered fixation interval of 3-4 s followed each trial in the event-related design. All events were presented in random order, counter-balanced for number size differences on the left or right side of the screen, and for number of left or right button presses.

**Figure 1.**
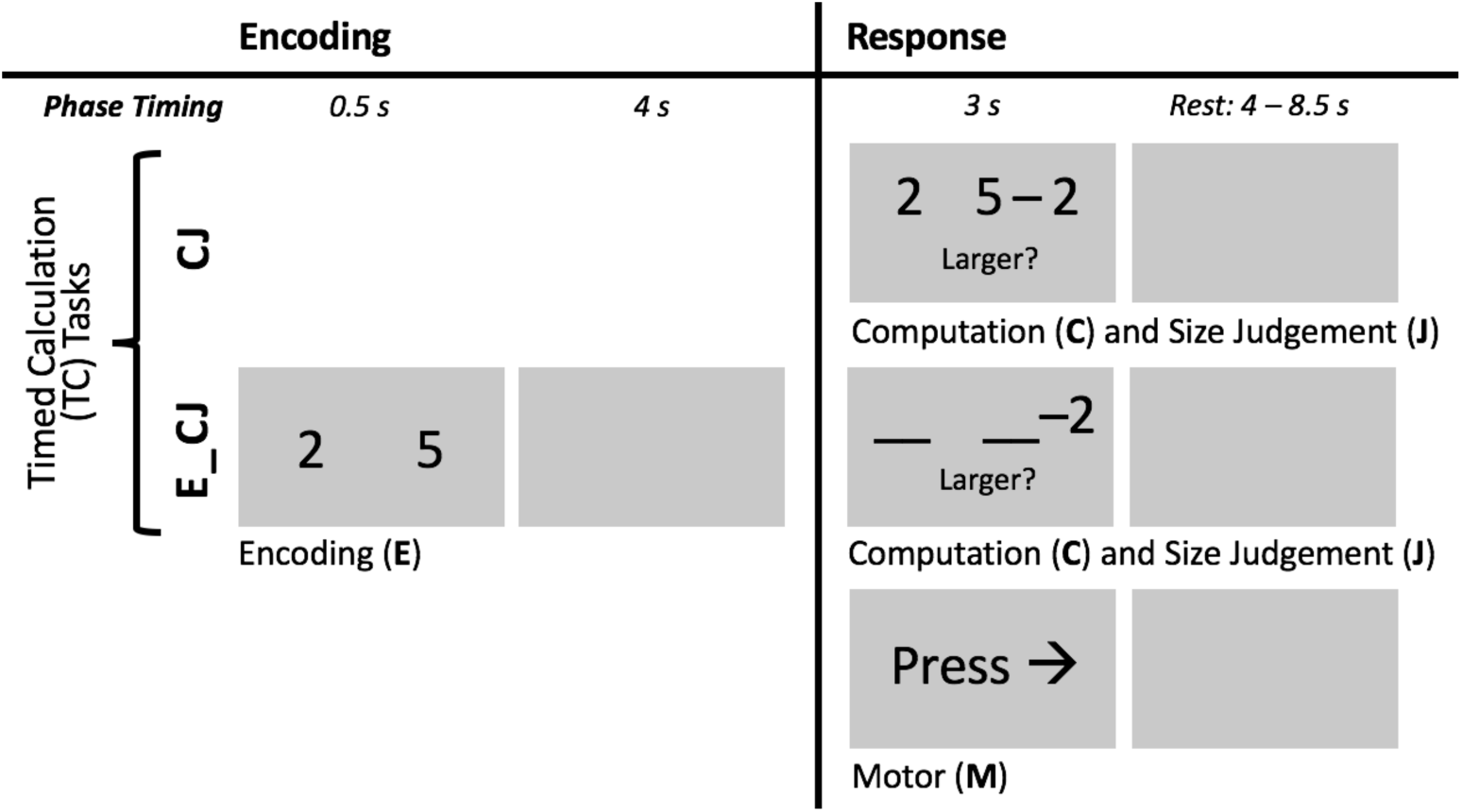
Math fluency-working memory (MF-WM) paradigm. In an event-related design with different trial types presented in pseudo-randomized order, subjects perform one of two math fluency (timed calculation) trials with response required within 3s. They performed arithmetic (subtraction of 2 or 3) from one of two single digits presented and indicated which of the resulting pair was larger (or smaller) as instructed (compute and numeric size judgement trials, CJ), or did the same after encoding (E) the two single digits in working memory. Control trials included pressing the left or right button as indicated (motor trials, M), and simple number size judgement (without arithmetic) after encoding of two single digits in working memory (working memory maintenance, not shown).

### Imaging Parameters

T2*-weighted echo planar imaging (EPI) images with BOLD contrast were obtained with a 3T MRI scanner (GEs, Milwaukee, WI) using a standard GE head coil (64 x 64 x 24 matrix with 3.75 x 3.75 x 6.0 mm spatial resolution, repetition time (TR) =2000 ms, echo time (TE) =28 ms, flip angle=90°, field of view (FOV) =24 x 24 cm) while participants performed the cognitive task. The first four scans were discarded to allow for signal saturation.

### fMRI Processing and Analyses

The functional imaging data were preprocessed and analyzed using the general linear model for event-related designs in statistical parametric mapping (SPM8, Wellcome Trust Centre for Neuroimaging, London, United Kingdom). The functional images were corrected for differences in acquisition time between slices for each whole-brain volume and realigned to correct for head movement. Six movement parameters (translation: x, y, z and rotation: roll, pitch, yaw) were included in the statistical model as covariates of no interest. The functional images were normalized to a standard EPI MNI template and then spatially smoothed using an isotropic Gaussian kernel of 8 mm full-width half-maximum.

A random-effects, event-related statistical analysis was performed at two levels. In the first level, the onsets and durations of each trial for each task condition were convolved with a canonical hemodynamic response function and modeled using a general linear model on an individual subject basis. The realignment parameters were included as additional regressors of no interest. Data were high-pass filtered at 1/128 Hz.

Random-effects analyses at the second (group) level in Experiments 1 and 2 (see below) were then conducted based on statistical parameter maps from each individual participant to allow population-level inference. Group level contrasts were thresholded at voxel-level p<0.001 and extent threshold to meet p<0.05 for cluster-level family-wise correction for multiple comparisons, unless otherwise stated.

We used Dynamic Causal Modeling, DCM (Friston et al., 2003) as implemented in SPM12 (DCM10) to examine how cortical and subcortical brain regions interacted during TC-related tasks in Experiments 1 and 2. Time series during TC were extracted from each of 6 regions-of-interest (ROI) in the IPS, AG, FFG, IFG, DLPFC and Str for each individual at a peak within 10mm of the group-level activation peak (at p<0.05 whole brain FWE corrected) and with p<0.05 at the individual subject level, as before (Kaplan C.M. et al., 2016). Specific ROI definitions are from Experiment 1 and detailed in the Results below. Deterministic DCM models comprising all possible pairings of these six ROIs (Nicholson et al., 2017) then enabled the estimation of the strength and direction of regional interactions to elucidate how regional neural activity and their interactions are influenced by cognitive inputs during TC, as well as how these neuronal effects are biophysically linked to form blood-oxygen-level-dependent signals (Friston et al., 2003). The resulting pairwise nodal connectivities describe how the cortical and subcortical brain regions interacted during TC tasks in Experiments 1 and 2. For each pair of regions, we constructed all 7 possible models of directed connectivity in one or both directions, with nodal inputs at one or both node pairs, which were then fitted to the observed data (Friston et al., 2011). Bayesian model averaging (BMA) was used to generate weighted task-related connectivity averages in each direction, for each pair of nodes, based on the posterior likelihood model fit (Penny et al., 2004; Stephan et al., 2007). Using these BMA results, we then examined task-related modulation of regional connectivity across participants, and group differences in Experiments 1 and 2. Permutation tests randomizing group labels over 1,000 iterations were performed across groups to detect differences in task-related effective connectivity without assumptions about probability distributions and to control for multiple comparisons (Nichols and Holmes, 2002).

## Results

### Experiment One

#### Demographic and Behavioral Results

Experiment 1 aimed to examine the neural correlates underlying TC in a comparison between otherwise healthy individuals with and without behavioral deficits in TC. This was examined in a cognitive paradigm in fMRI, where subjects performed simple arithmetic and number size comparisons on single digit integers (Tan et al., 2007) (Figure 1). We defined math fluency deficit (MFD, n=34, 17 males) as those individuals who performed less accurately on TC tasks, but not working memory maintenance (accuracy: TC < 0.5, working memory maintenance ≥ 0.8). The control (CON) group (n = 34, 17 males) were participants who performed well in both tasks (working memory maintenance and TC accuracy ≥ 0.8, mean accuracy = 0.92). Both groups were similar for age, gender, and years of education. There were group differences in IQ (Wechsler Adult Intelligence Scale, WAIS) stemming from the arithmetic component of WAIS, providing additional evidence of math dysfunction in MFD vs CON individuals (Table 1).

**Table 1.**
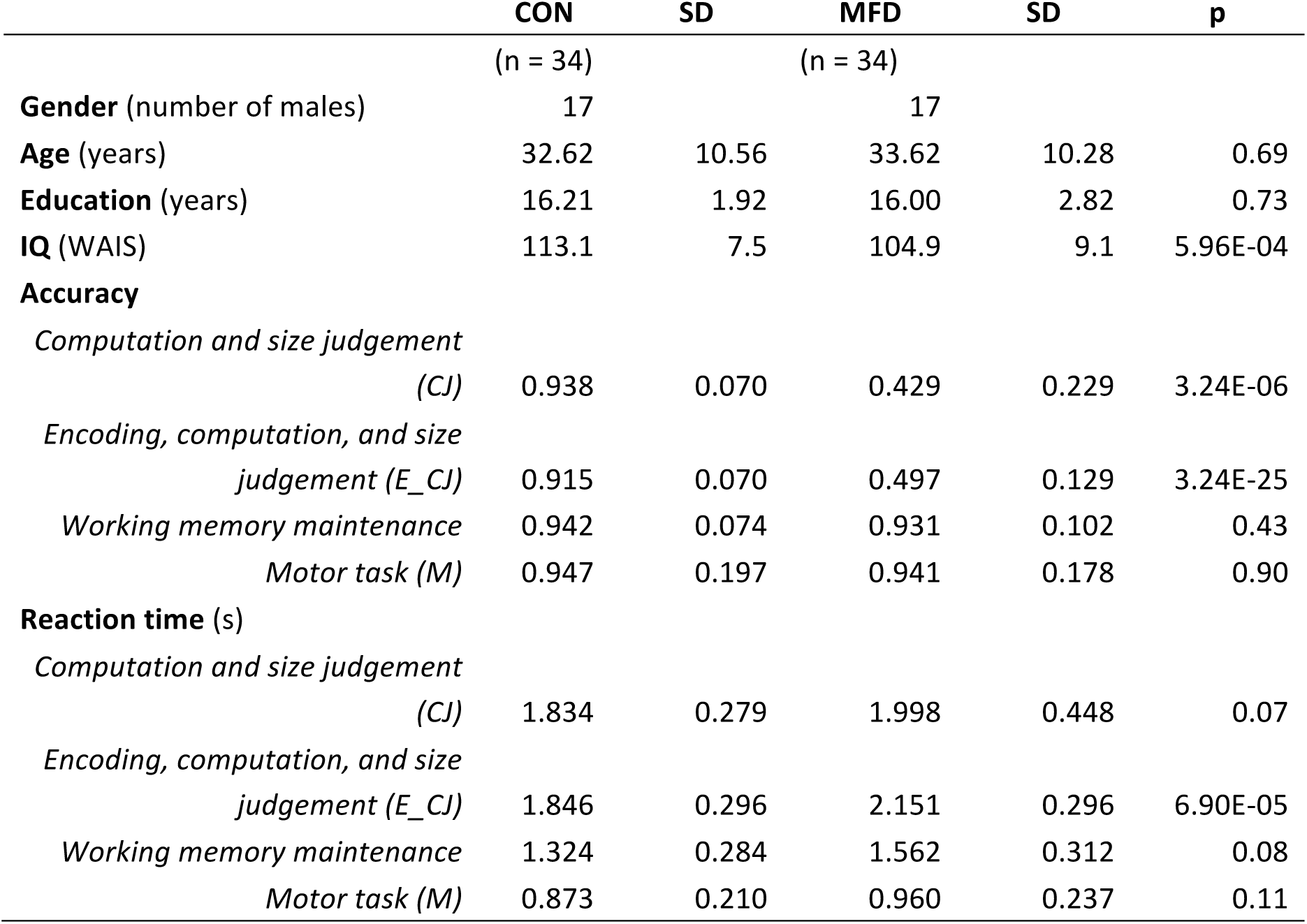
Demographics and Behavioral Performance for CON and MFD Groups (Exp. 1).

While MFD participants were, by definition, less accurate at arithmetic tasks, the CON and MFD groups did not differ in the accuracy of working memory maintenance and motor control tasks (Table 1). However, MFD participants were slower at all number-related tasks, though not for the motor response task.

#### fMRI Results

One-sample t-test activation during TC of each CON and MFD group are in Supplementary Tables 1 and 2. During TC, CON participants showed stronger activation than MFD in the right IPS (48 −44 50, T=5.01, k_E_=128, voxel-wise p<0.001 and cluster-family-wise error, FWE corrected p<0.05, Figure 2a; other regions differentially engaged are in Supplementary Table 3). There were no regions where MFD participants had relatively increased activation.

**Figure 2.**
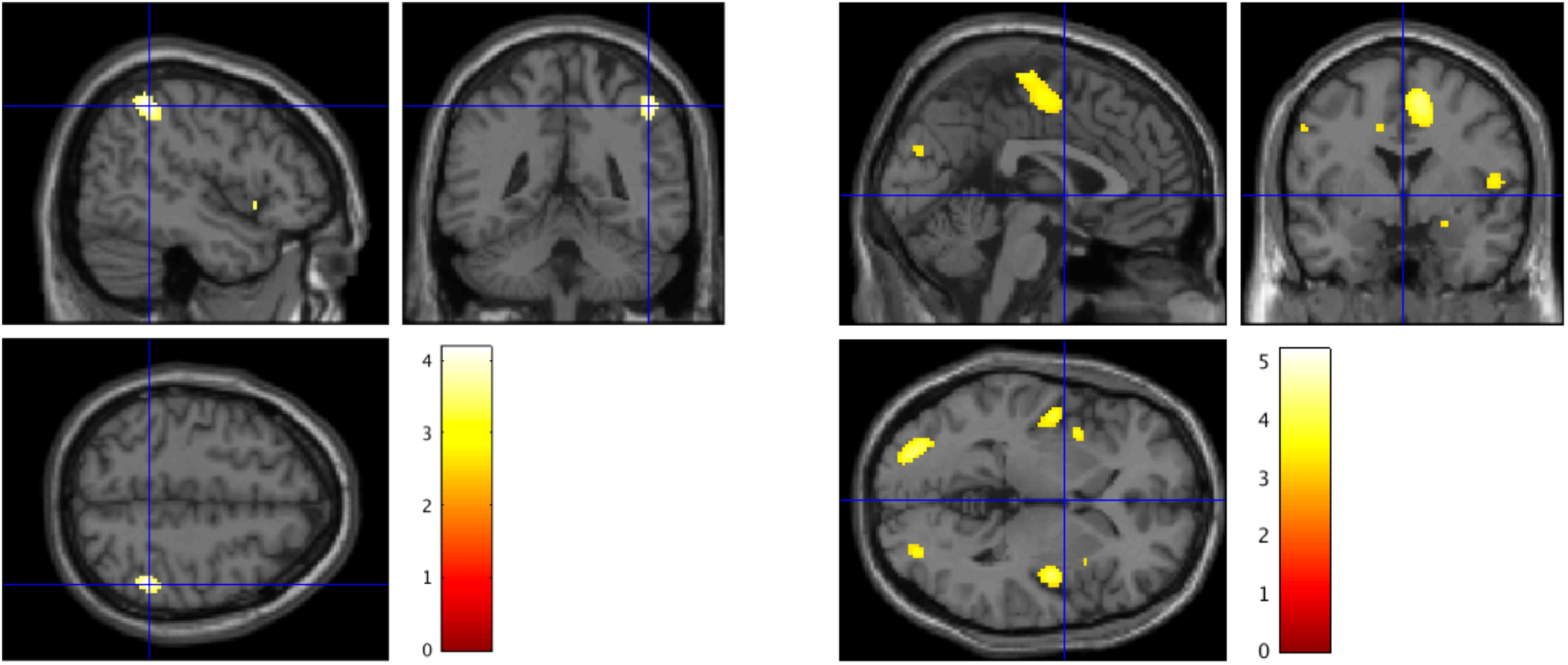
Task-related activation contrasts. (a) During the math fluency-working memory paradigm task (CJ, ECJ), in Experiment 1, the CON group (N=34) engaged the right IPS [48-44 50] more strongly compared to the MFD group (N=34), p<0.05 cluster-FWE corrected. (b) In Experiment 2, MF-WM paradigm task contrast engaged increased activation in ‘compensated’ low-IPS (N=50) compared to high-IPS (N=50), at fusiform gyrus, inferior frontal gyrus (and not shown, angular gyrus and putamen, p<0.05 whole brain FWE-corrected).

#### Effective Connectivity Results

Using Dynamic Causal Modeling (DCM) in SPM12, we examined how effective connectivity differed between MFD and CON groups. Regions of interest (ROIs) were modeled pair-wise in DCM (Nicholson et al., 2017) (see Methods), and were defined as follows. The IPS peak (48 −44 50, above) that was differentially engaged across MFD vs CON was defined as one ROI. Our goal was to examine how connectivity between this IPS ROI and other ROIs from the literature that have functional and structural connections with each other (Hearne et al., 2015; Horwitz et al., 1998; Jung et al., 2017; Uddin et al., 2010), and are engaged in numerical cognition and TC (Arsalidou and Taylor, 2011; Dehaene et al., 2004), may differ across MFD and CON groups. The other ROIs included the angular gyrus (AG, 34 −60 30), fusiform gyrus (FFG, 34 −80 −6), inferior frontal gyrus (IFG, 36 24 4), dorsolateral prefrontal cortex (DLPFC, 46 30 26) and striatum (Str, 18 10 0). All these subsequent ROIs were defined from the highest peaks in the MFD and CON groups in the Pickatlas defined regions (Maldjian et al., 2003) that were robustly activated (p<0.05 whole brain family-wise error corrected for multiple comparisons) in each MFD and CON group (Supplementary Tables 1 and 2), and were not differentially engaged between MFD and CON. The same ROIs were also used in Experiment 2 (below), which comprised an independent sample of healthy subjects without deficits in TC.

In Experiment 1, bidirectional effective connectivity from Bayesian Model Averaging across all 7 possible models in each pair of nodes over the 6 regions-of-interest (IPS, AG, FFG, IFG, DLPFC and Str) were mostly significant in one-sample t-tests in each of the CON and MFD groups (Supplementary Tables 6 and 7). However, the MFD group had relatively reduced connectivity between IPS and Str (IPS-to-Str: permutation p=0.03, Str-to-IPS: permutation p=1.89E-03), and between DLPFC and Str (DLPFC-to-Str: permutation p=9.66E-03, Str-to-DLPFC: permutation p<0.001) (Figure 3a, Figure 4a). In MFD, Str connectivity was also decreased to IFG (permutation p<0.01) and FFG (permutation p=5.28E-03), and there was decreased connectivity to DLPFC from AG (permutation p<0.04) and from IPS (permutation p=0.04). Significant decreases in effective connectivity in MFD also included IFG-to-IPS (permutation p=6.42E-03), and DLPFC-to-IFG (permutation p=3.85E-03). There was relatively increased DLPFC-to-AG (permutation p<0.02) connectivity in MFD. We then explored how some of these changes may be compensated for in Experiment 2, positing that increased engagement of pathways bypassing decreased IPS activation and connectivity may be compensatory if subjects maintained high performance despite reduced IPS-related engagement.

**Figure 3:**
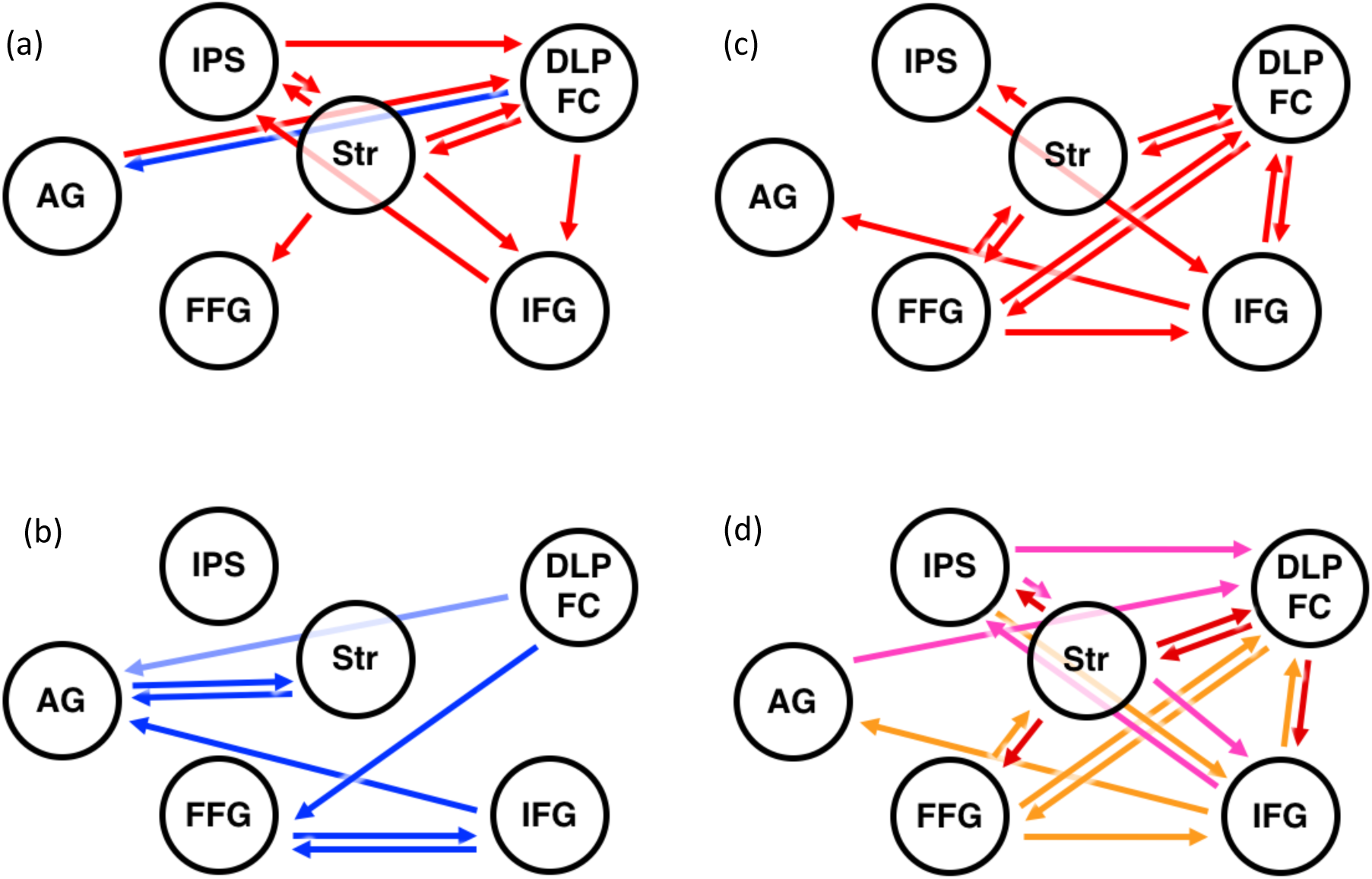
DCM connectivity contrasts during MF. (a) Exp. 1: Network model of Bayesian model averaging values of effective connectivity in MFD (N=34) relative to CON (N=34) group. Blue = effective connectivity increase (MFD>CON); red = decrease (MFD<CON). Notably, MFD had decreased effective connectivity in DLPFC input (3 connections) and Str output (4 connections). (b) Exp. 2: Network model of significant increases in Bayesian model averaging values of effective connectivity in ‘compensated’ low-IPS (N=50) relative to high-IPS group (N=50). No significant differences occurred in IPS connectivity. Blue = significant effective connectivity increase (low-IPS>high-IPS). (c) Differences in node input effective connectivity for MFD (N=34) relative to ‘compensated’ low-IPS (N=50). Red = significant effective connectivity decrease (MFD<low-IPS). (d) Overlap in decreases between MFD relative to CON (Figure 3a), and MFD relative to compensated low-IPS comparison (Figure 3c). Magenta: MFD < CON only; orange: MFD < low-IPS only; dark red: MFD < CON and MFD < low-IPS. See Supplementary Tables 6-8.

**Figure 4:**
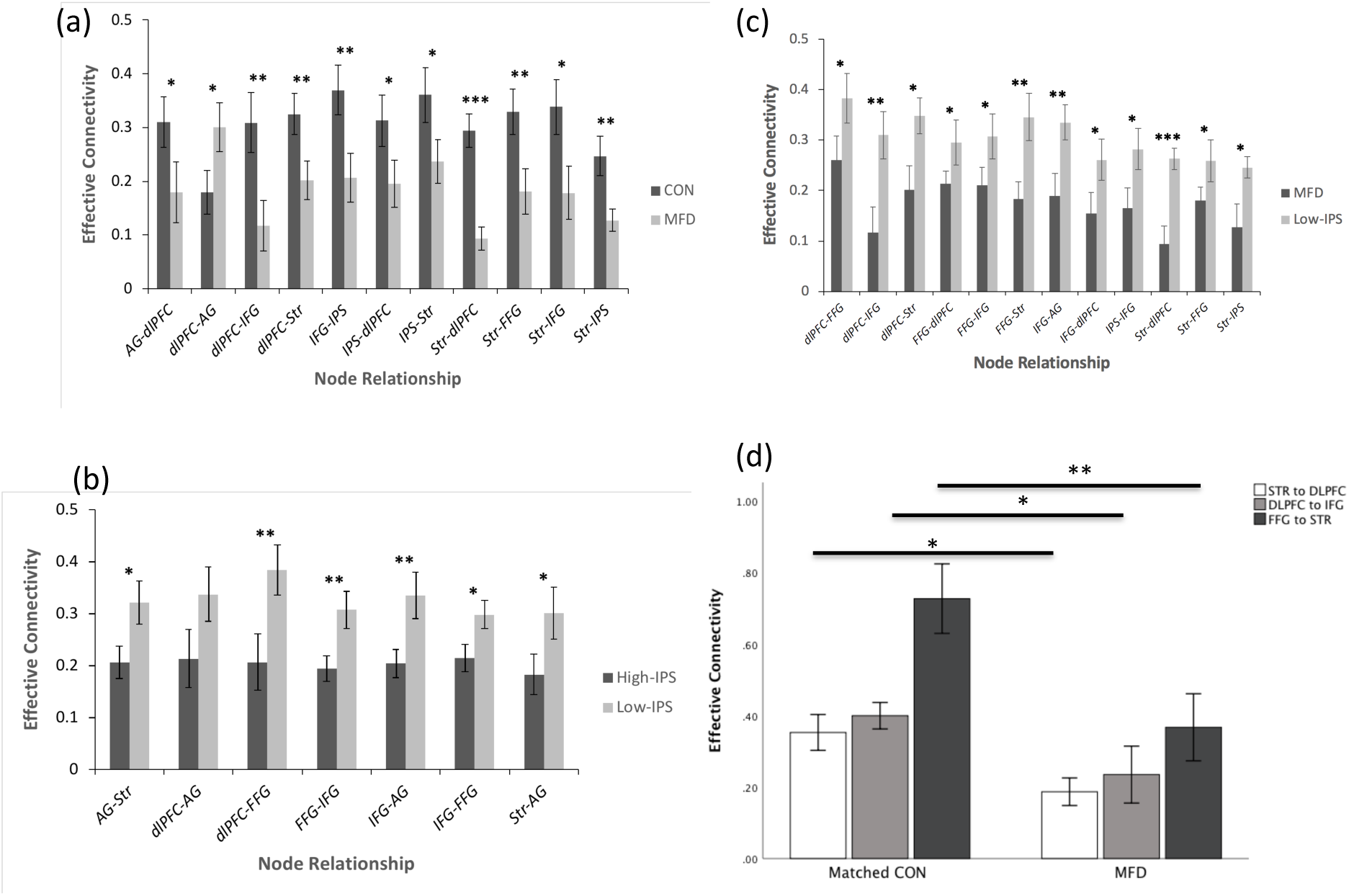
Effective connectivity across groups. (a) Significantly different effective connectivity (labeled as ‘from’ region A ‘to’ B) across CON (N=34) and MFD (N=34, Exp 1), (b) across ‘compensated’ high-IPS (N=50) and low-IPS (N=50) groups (Exp 2), and (c) across ‘compensated’ low-IPS (N=50) and MFD (N=34) groups. (d) Significantly different effective connectivity across controls matched using support vector regression (Matched CON N=46) to MFD (N=34) for DLPFC, Str and IPS connectivity vs. MFD. Permutation p-values *p<0.05, **p<0.005, ***p<0.001.

### Experiment 2

#### Behavioral Results

Experiment 2 examined the neural correlates of IPS deficits and potential compensatory function during TC in an independent group of healthy individuals who had no behavioral deficits in TC (n = 100, 45 males; accuracy ≥ 0.8 for TC). We first divided the subjects into two equal subgroups of relatively high and low IPS ROI activity (above mean activation, n=50, 26 males; below mean activation, n=50, 19 males). The groups engaging relatively high (N=50) and low IPS activation (N=50) did not otherwise differ in age, education, IQ, or TC accuracy and response time (Table 2).

**Table 2.**
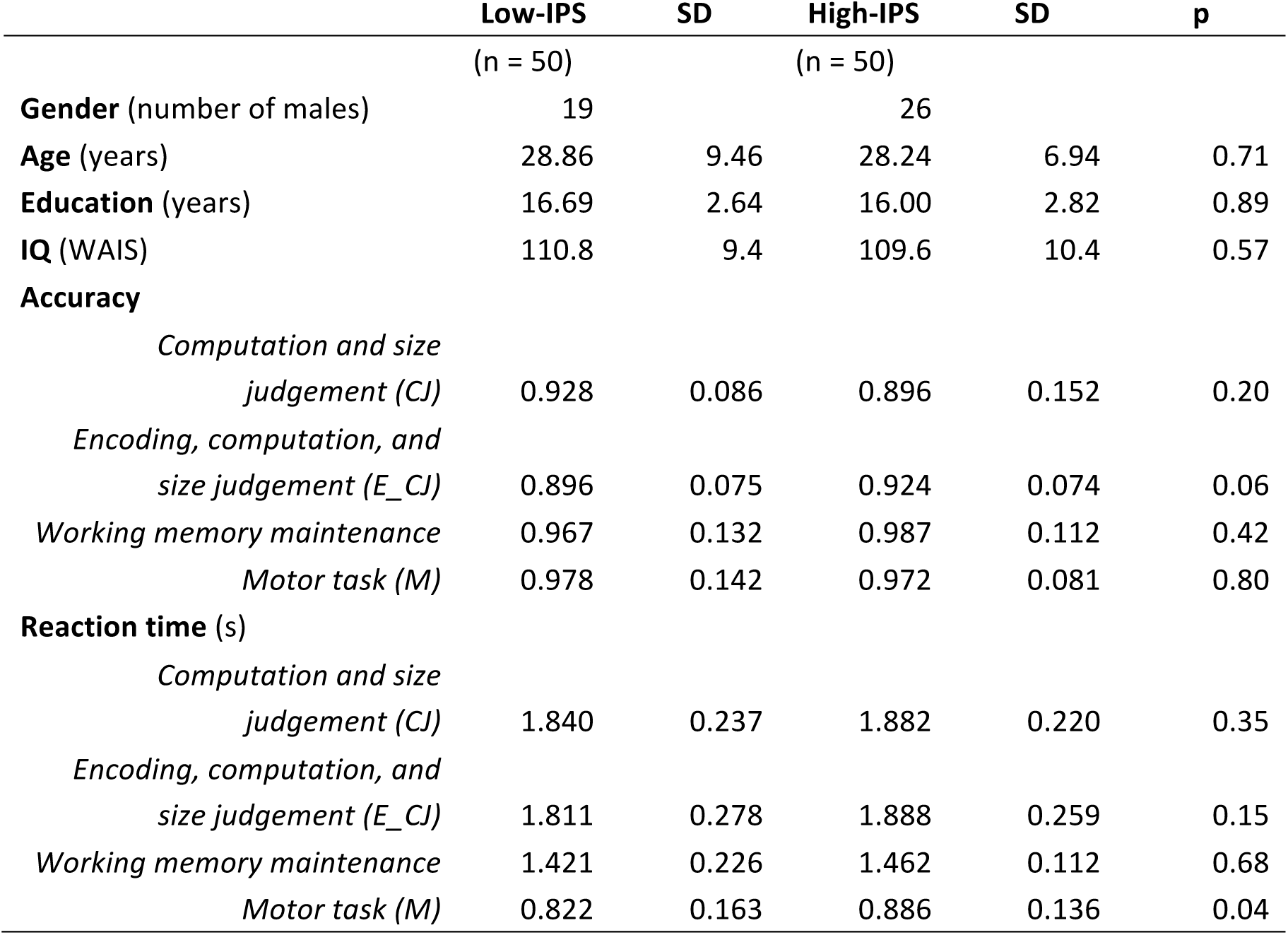
Demographics and Behavioral Performance for Low-IPS and High-IPS Groups (Exp. 2).

#### fMRI Results

Both groups, combined, robustly engaged IPS, AG, FFG, Str, IFG and DLPFC during TC (Supplementary Table 4). There was, by definition, significantly stronger IPS activation in the high-IPS group at TC (48 −42 48, T=8.28, k_E_=616, p<0.05 cluster FWE corrected). Relative to the high-IPS group, the equally performing low-IPS group, however, also showed stronger engagement of the AG (40 −58 28, T=3.52, k_E_=48), FFG (32 −84 −2, T=3.44, k_E_=45), and IFG (36 10 −10, T=3.21, k_E_>100) (Figure 2b). Both groups had similarly robust activation at the DLPFC (46 30 26, T>18, k_E_>100) and Str (14 6 0, T>13, k_E_>50). Differences across groups in other brain regions are detailed in Supplementary Table 5.

#### Effective Connectivity Results

All pairwise effective connectivities between the 6 IPS, AG, FFG, Str, IFG and DLPFC ROIs, defined from Experiment 1, and applied to the independent sample here in Experiment 2, significantly differed from zero in the combined high and low IPS groups (Supplementary Table 8). Across low-IPS vs. high-IPS groups, there were relatively increased ‘compensatory’ excitatory effective connectivities in the low vs high-IPS group for AG-Str (AG-to-Str: permutation p=0.01, Str-to-AG: permutation p=0.03) and FFG-IFG (FFG-to-IFG: permutation p=3.93E-03, IFG-to-FFG: permutation p=0.02) (Figure 3b, Figure 4b). We also found relatively increased excitatory effective connectivity in the low-IPS group relative to the high-IPS group from DLPFC-to-FFG (permutation p=3.70E-03), and IFG-to-AG (permutation p=5.26E-03). There was no decreased connectivity between the IPS and DLPFC, Str and IFG in the low vs high IPS group. Thus, it would appear that despite reduced engagement of IPS, maintained performance in TC was associated with a network of increased ‘compensatory’ connectivity at AG, FFG, IFG, Str and DLPFC, and maintained effective connectivity to and from IPS.

We next examined connectivity across the poorly performing MFD group vs. the ‘compensated’ low IPS activation group, to further define connectivity bypassing IPS that could be relevant in behavioral compensation (Figure 3c, 4c). Connectivity was significantly increased in the low-IPS group relative to MFD at DLPFC-FFG (DLPFC-to-FFG: permutation p=0.04, FFG-to-DLPFC: permutation p=0.04), DLPFC-IFG (DLPFC-to-IFG: permutation p=2.90E-03, IFG-to-DLPFC: permutation p=0.03), DLPFC-Str (DLPFC-to-Str: permutation p=0.01, Str-to-DLPFC: permutation p<0.001), FFG-Str (FFG-to-Str: permutation p=1.01E-03, Str-to-FFG: permutation p=0.04). Connectivity increases in the low-IPS group also included that of IFG from FFG (permutation p=0.04) and from IPS (permutation p=0.02), IFG-to-AG (permutation p=6.43E-03), and Str-to-IPS (permutation p=0.02). On the other hand, while connectivity between IPS and DLPFC was reduced in MFD vs. CON (Exp 1), they were not different in MFD vs. low IP group (Figure 3d, magenta connections). Compensated subjects with reduced IPS activation thus appeared to have increased engagement of ventral cortical networks in processing TC, which overlapped in comparisons between low IPS vs high IPS groups, and between low IPS vs MFD groups (Figure 3d).

We then examined a different selection of compensated subjects based on IPS connectivity instead of IPS activation. Specifically, we examined those with reduced dorsal corticostriatal connectivity patterns in IPS, DLPFC, and Str, similar to that occurring in MFD (Exp 1). We examined how reduced connectivity in these dorsal IPS-related networks, well known to be important for TC (Arsalidou and Taylor, 2011; Dehaene et al., 2004) may be bypassed in maintaining performance. Here, we created a support-vector regression model differentiating MFD from CON using DCM connectivity in DLPFC, IPS and Str from subject data in Experiment 1. We then applied this model to the high-performing subjects in Experiment 2, to identify a subgroup of 46 subjects that had similar reductions in connectivity patterns in these brain regions, but ostensibly with compensated behavior. In this subgroup of higher performing individuals, there were, indeed, no significant differences in the IPS-to-DLPFC and IPS-to-Str connectivities with that in MFD. However, relative to MFD, there remained a network of putatively increased compensatory connectivity from FFA to Str, from Str to DLPFC, and from DLPFC to IFG in the compensated group of high performing individuals (p<0.05 permutation test, Figure 4d).

## Discussion

We examined independent samples of otherwise healthy individuals with differing timed calculation performance that suggest individual variation in apparent re-balancing of neural connectivity to maintain TC performance. In Experiment 1, CON participants engaged the right IPS more strongly than those who performed TC poorly, consistent with the known importance of the IPS in numerical processing (Arsalidou and Taylor, 2011; Cohen Kadosh and Walsh, 2009; Dehaene et al., 2003; Eger et al., 2003; Pinel et al., 2004; Venkatraman et al., 2005). Across an extended network of brain regions previously implicated in numerical cognition and TC (Arsalidou and Taylor, 2011; Dehaene et al., 2004), MFD was associated with reduced effective connectivity from IPS to DLPFC and to Str, as well as reduced effective connectivity across cortical regions in the DLPFC, FFG, IFG, IPS and striatum (Figure 3a).

In Experiment 2, we studied an independent sample of individuals who performed equally well in TC but varied in IPS activity, with the goal of exploring potential network variation that may compensate for reduced IPS engagement. We found that individuals with relatively low-IPS engagement but preserved TC performance had increased engagement of the DLPFC, AG, FFG, IFG, and Str. On one hand in Experiment 1, MFD was associated with reduced effective connectivity from IPS to DLPFC and to Str, as well as reduced Str effective connectivity to cortical regions in the DLPFC, FFG, IFG, and IPS (Figure 3a). On the other hand, in Experiment 2, subjects with maintained TC performance but relatively reduced IPS engagement had increased Str connectivity to DLPFC, IPS and AG (Figure 3b). A comparison across MFD and compensated low-IPS groups further supported the Str effect – the poorly performing MFD group had decreased Str connectivity with DLPFC, FFG, and IPS relative to high-performing subjects with similarly low IPS engagement and/ or connectivity (Figure 3c, 4c). These findings may thus reflect the role Str plays in utilizing dopaminergic engagement to effectively gate new information during cortical processing needed in numerical calculations (Landau et al., 2009; O’Reilly, 2006) (Arsalidou and Taylor, 2011; Dehaene et al., 2004) and cortical-striatal connectivity in engaging working memory and attentional processing integral in TC performance (Askenazi and Henik, 2010; Corbetta and Shulman, 2002; Frank et al., 2001; Marklund and Persson, 2012).

Moreover, we found significantly increased effective connectivity between FFG, Str, DLPFC and IFG in the compensated low IPS vs. high IPS groups (Figure 3b, 4b), which also occurred across the low IPS vs. MFD groups (Figure 3c, 4c), and in high performing individuals with similar deficits in IPS-Str-DLPFC connectivity as MFD (Figure 4d). It appears these ventral cortical connectivity may help maintain performance despite reduced IPS engagement and connectivity, possibly by re-balancing some of the reduced IPS activation and connectivity patterns through strengthened activation and effective connectivity, resulting in compensated performance.

Of the two distinct paradigmatic visual information processing streams, i.e., the dorsal ‘where’ and ventral ‘what’ streams (Ungerleider and Haxby, 1994), the dorsal visual stream includes cortical regions such as the IPS and DLPFC (Rubinsten and Henik, 2009; Ungerleider and Haxby, 1994). The ventral visual stream and interactions between the AG, FFG, and IFG has recently been suggested to be engaged during working memory numerical tasks, retrieval of simple arithmetic facts, and linguistic and symbolic representation of numerical concepts (Rubinsten and Henik, 2009; Wilson and Dehaene, 2007). Our findings of increased connectivity between FFG, Str, and IFG in compensating TC performance suggest that there may exist interactions between these two paradigmatic networks in maintaining math fluency. MFD had decreased connectivity in the dorsal network (Figure 3c), while the compensated groups appeared to have increased ventral stream connectivity, and interactions between these streams through the DLPFC (Figure 3d). It may be plausible that at least in some individuals with reduced IPS engagement and connectivity, mechanisms of neuroplasticity or learning could have been involved, during learning of arithmetic in childhood, in maintaining TC performance through differential engagement of these ventral cortical networks. If so, strategies to enhance the function of these networks may be useful.

It may be possible that differing cognitive strategies in the compensated low-IPS individuals may contribute to the varying patterns of cortical engagement we found. While working memory is needed to hold and manipulate numerical information, retrieval of arithmetic facts may also contribute to math fluency (Locuniak and Jordan, 2008). Indeed, the low-IPS group had significantly increased AG activation and IFG-to AG connectivity compared with the high-IPS and MFD groups. It has been suggested that children with poor math fluency rely more strongly on fact retrieval to complete arithmetic problems engaging AG (De Smedt et al., 2011). Thus, one possibility is that the low-IPS group used fact retrieval strategies for less-effortful, quicker problem solving with small numbers during the MF- working memory paradigm, engaging the AG (Dehaene et al., 2003; Grabner et al., 2009; Stanescu-Cosson et al., 2000). Not only has fact retrieval training been shown to increase activation in the AG, but activation in the IPS may decrease as well (Delazer et al., 2003; Grabner et al., 2009). However, the timed calculation tasks in our MF paradigm comprised of small-number subtraction. Number sense is thought to be more reliably engaged in subtraction operations engaging the IPS, while addition is more dependent on language-based fact retrieval in the angular gyrus (De Smedt et al., 2011; Dehaene et al., 2004). We are also unaware of any literature suggesting roles of FFG in fact retrieval. While we cannot rule out that AG-associated arithmetic fact retrieval has not also occurred, we posit that it may be more likely a distinct compensatory mechanism involving FFG and associated ventral networks is also engaged in maintained math fluency performance, distinct from arithmetic fact retrieval. Indeed, the engagement of ventral high-level visual areas have also recently been found during math tasks in individuals with congenital blindness, suggesting plasticity in these functions (Kanjlia et al., 2019), that we posit may also occur to beyond individuals with visual impairment.

In conclusion, we found that calculation difficulty is associated with well-established reduced IPS engagement, and a pattern of reduced Str to cortical (IPS, DLPFC, FFG, and IFG) effective connectivity. However, subsets of individuals with reduced IPS engagement and connectivity could maintain MF performance. This occurred through increased effective connectivity at the DLPFC, FFG, IFG, and Str. The genetic and environmental contributions to these variations in brain network engagement remain to be understood. Our results reinforce notions that a distributed network of brain regions including AG, FFG, and IFG process TC (Arsalidou and Taylor, 2011; Dehaene et al., 2004) and further suggests that there could be significant inter-individual variability and rebalancing in the engagement of these FFG-related ventral cortical networks in TC when IPS-related network connectivity are reduced. The extent to which these individual variations could be leveraged, for example, in personalized remediation strategies for dyscalculia remains to be determined in future work.

## Acknowledgements

This work was funded by the National Institute of Mental Health Intramural Research Program (Daniel R Weinberger, KFB), Lieber Institute for Brain Development, and National Institute of Mental Health grant R01MH101053 (HYT).

