## Supplementary materials for "Dysfunctional and compensatory brain networks underlying math fluency"

**Supplementary Table 1**
One-sample t-test activation data (SPM12; thresholded at p<0.05 FWE corrected) of control (CON, N=34) group during timed calculation (TC).

| Region | BA | Coordinates | k | z |
|---|---|---|---|---|
| L medial frontal gyrus | 6 | -2 -2 58 | 86009 | > 20 |
| R superior frontal gyrus | 6 | 6 8 52 | | > 20 |
| R middle frontal gyrus | 9/46 | 52 26 26 | | >20 |
| R inferior frontal gyrus | 44 | 34 22 -2 | | >20 |
| L inferior parietal lobule | 40 | -48 -38 46 | | > 20 |
| L inferior frontal gyrus | 9 | -52 4 28 | | > 20 |
| R inferior parietal lobule | 40 | 46 -46 48 | | > 20 |
| R angular gyrus | | 36 -58 30 | | >20 |
| R fusiform gyrus | | 36 -90 2 | | >20 |
| R cingulate gyrus | 23 | 8 -30 26 | 668 | 7.31 |
| R putamen | | 14 8 2 | | 10.9 |
| L posterior cingulate | 23 | -4 -34 24 | | 6.56 |

BA, Brodmann area; k, cluster size; z, z-statistic; R, right; L, left.

**Supplementary Table 2**
One-sample t-test activation data (SPM12; thresholded at p<0.05 FWE corrected) of math fluency dysfunction (MFD, N=34) group during timed calculation (TC).

| Region | BA | Coordinates | k | z |
|---|---|---|---|---|
| L middle occipital gyrus | 19 | -34 -88 16 | 2380 | > 20 |
| R middle frontal gyrus | 9/46 | 44 28 22 | | 7.1 |
| R inferior frontal gyrus | 44 | 34 24 4 | | 8.7 |
| R angular gyrus | | 34 -60 28 | | 4.8 |
| R fusiform gyrus | | 38 -86 -8 | | 7.8 |
| L thalamus | | -10 -20 2 | | 6.25 |
| L putamen | | -20 8 -2 | | 5.89 |
| R putamen | | 18 10 -4 | | 5.86 |
| R cerebellum | | 22 -68 -30 | 14 | 5.2 |
| R cerebellum | | 8 -82 -30 | 52 | 5.13 |
| L cerebellum | | -8 -82 -30 | | 4.81 |
| R cerebellum | | 2 -58 -40 | 29 | 4.92 |
| L middle temporal gyrus | 39 | -54 -58 10 | 30 | 4.54 |
| R posterior cingulate | 23 | 6 -34 24 | 7 | 4.5 |
| Medulla | | 2 -36 -44 | 3 | 4.35 |

BA, Brodmann area; k, cluster size; z, z-statistic; R, right; L, left.

**Supplementary Table 3**
Two-sample t-test activation data (SPM12; thresholded at $p<0.05$ FWE corrected) of control (CON, N=34) – math fluency dysfunction (MFD, N=34) groups during timed calculation (TC).

| Region | BA | Coordinates | k | z |
|---|---|---|---|---|
| L insula | | -36 10 2 | 124 | 5.5 |
| R inferior frontal gyrus | 47 | 34 24 -16 | 60 | 5.18 |
| R cerebellum | | 22 -60 -30 | 80 | 5.12 |
| R cerebellum | | 12 -62 -34 | | 4.52 |
| R inferior frontal gyrus | 47 | 44 16 -4 | 89 | 5.06 |
| L superior frontal gyrus | 6 | -4 10 60 | 139 | 5.06 |
| R intraparietal sulcus | 40 | 48 -44 50 | 128 | 5.01 |
| R cerebellum | | 8 -68 -6 | 43 | 4.66 |
| L precentral gyrus | 4 | -30 -28 62 | 31 | 4.55 |
| R precuneus | 7 | 24 -78 44 | 12 | 4.47 |
| L cerebellum | | -24 -34 -32 | 1 | 4.43 |
| R cerebellum | | 16 -62 -6 | 3 | 4.31 |
| L inferior parietal lobule | 40 | -48 -40 42 | 1 | 4.3 |

BA, Brodmann area; k, cluster size; z, z-statistic; R, right; L, left.

**Supplementary Table 4**
One-sample t-test activation data (SPM12; thresholded at $p<0.05$ FWE-corrected) during timed calculation (TC) in Exp 2 (N=100)

| Region | BA | Coordinates | k | z |
|---|---|---|---|---|
| L inferior frontal gyrus | 9 | -46 4 28 | 100980 | > 20 |
| L superior parietal lobule | 7 | -26 -72 44 | | > 20 |
| L superior frontal gyrus | 6 | -4 6 54 | | > 20 |
| L inferior parietal lobule | 40 | -34 -54 44 | | > 20 |
| R superior frontal gyrus | 6 | 6 8 54 | | > 20 |
| L inferior parietal lobule | 40 | -40 -50 46 | | > 20 |
| R inferior frontal gyrus | 9 | 46 8 26 | | > 20 |
| R superior parietal lobule | 7 | 30 -70 44 | | > 20 |
| L inferior frontal gyrus | 47 | -32 22 -4 | | > 20 |
| R superior parietal lobule | 7 | 36 -64 46 | | > 20 |
| L precentral gyrus | 4 | -38 -14 58 | | > 20 |
| R inferior frontal gyrus | 47 | 34 24 -2 | | > 20 |
| R inferior parietal lobule | 40 | 40 -52 44 | | > 20 |
| R dorsolateral prefrontal cortex | 9 | 46 30 26 | | > 20 |
| R globus pallidus | | 14 6 0 | | > 20 |
| R sub-gyral temporal lobe | | 32 -2 -26 | 3 | 4.74 |

BA, Brodmann area; k, cluster size; z, z-statistic; R, right; L, left.

**Supplementary Table 5**
Two-sample t-test activation data (SPM12; thresholded at p<0.05 FWE-corrected) of low-IPS (N=50) vs high-IPS (N=50) groups during timed calculation (TC).

| Region | BA | Coordinates | k | z |
|---|---|---|---|---|
| R cingulate gyrus | 24 | 12 -6 46 | 1418 | 6.21 |
| R cuneus | 19 | 6 -84 26 | 1737 | 5.81 |
| R insula | 40 | 48 -22 14 | 534 | 5.78 |
| R precentral gyrus | 44 | 46 2 6 | 1129 | 5.65 |
| R inferior frontal gyrus | 13 | 36 10 -10 | | 5.44 |
| R angular gyrus | 39 | 40 -58 26 | 184 | 5.56 |
| L sub-gyral temporal lobe | | -30 -60 28 | 412 | 5.54 |
| L lingual gyrus | | -26 -88 -2 | 149 | 5.2 |
| L superior temporal gyrus | 22 | -48 -4 4 | 774 | 5.11 |
| L inferior parietal lobule | 40 | -66 -34 22 | 174 | 5.07 |
| L sub-gyral parietal lobe | | -30 -30 38 | 192 | 4.92 |
| L precentral gyrus | | -54 -8 38 | 47 | 4.9 |
| L superior temporal gyrus | 38 | -52 6 -12 | 33 | 4.78 |
| R cerebellum | | 6 -80 -10 | 160 | 4.78 |
| R fusiform gyrus* | 18 | 32 -84 -2 | | 3.75 |

BA, Brodmann area; k, cluster size; z, z-statistic; R, right; L, left.
*Did not survive FWE correction

**Supplementary Table 6**
One-sample t-tests of dynamic causal modeling (DCM) and Bayesian model averaging (BMA) node relationship data for control (CON, N=34) group during timed calculation (TC) in Exp 1.

| Node Relationships | Mean | SD | P-values | T-values |
|---|---|---|---|---|
| AG-DLPFC | 0.34 | 0.14 | < 0.001 | 10.34 |
| DLPFC-AG | 0.24 | 0.12 | < 0.001 | 7.48 |
| AG-FFG | 0.37 | 0.14 | < 0.001 | 7.39 |
| FFG-AG | 0.25 | 0.13 | < 0.001 | 8.70 |
| AG-IFG | 0.28 | 0.14 | < 0.001 | 7.88 |
| IFG-AG | 0.26 | 0.14 | < 0.001 | 8.94 |
| AG-IPS | 0.38 | 0.14 | < 0.001 | 9.10 |
| IPS-AG | 0.34 | 0.13 | < 0.001 | 6.89 |
| AG-Str | 0.30 | 0.14 | < 0.001 | 8.46 |
| Str-AG | 0.30 | 0.15 | < 0.001 | 6.61 |
| DLPFC-FFG | 0.39 | 0.14 | < 0.001 | 8.43 |
| FFG-DLPFC | 0.30 | 0.13 | < 0.001 | 8.11 |
| DLPFC-IFG | 0.31 | 0.14 | < 0.001 | 9.68 |
| IFG-DLPFC | 0.26 | 0.16 | < 0.001 | 5.46 |
| DLPFC-IPS | 0.50 | 0.14 | < 0.001 | 6.46 |

| | | | | |
|---|---|---|---|---|
| IPS-DLPFC | 0.32 | 0.12 | < 0.001 | 5.78 |
| DLPFC-Str | 0.27 | 0.14 | < 0.001 | 6.12 |
| Str-DLPFC | 0.42 | 0.16 | < 0.001 | 7.26 |
| FFG-IFG | 0.29 | 0.15 | < 0.001 | 8.58 |
| IFG-FFG | 0.28 | 0.15 | < 0.001 | 6.11 |
| FFG-IPS | 0.29 | 0.12 | < 0.001 | 6.11 |
| IPS-FFG | 0.40 | 0.14 | < 0.001 | 11.93 |
| FFG-Str | 0.24 | 0.15 | < 0.001 | 9.08 |
| Str-FFG | 0.43 | 0.15 | < 0.001 | 9.40 |
| IFG-IPS | 0.54 | 0.17 | < 0.001 | 6.62 |
| IPS-IFG | 0.18 | 0.12 | < 0.001 | 8.66 |
| IFG-Str | 0.23 | 0.15 | < 0.001 | 7.94 |
| Str-IFG | 0.53 | 0.17 | < 0.001 | 8.09 |
| IPS-Str | 0.21 | 0.12 | < 0.001 | 6.46 |
| Str-IPS | 0.43 | 0.16 | < 0.001 | 6.44 |

Node relationship labels follow “from - to” directionality; all nodes are right hemisphere, comprising angular gyrus (AG), dorsolateral prefrontal cortex (DLPFC), fusiform gyrus (FFG), inferior frontal gyrus (IFG), intraparietal sulcus (IPS), and striatum (Str).

**Supplementary Table 7**

One-sample t-tests of dynamic causal modeling (DCM) and Bayesian model averaging (BMA) node relationship data for math fluency dysfunction (MFD, N=34) group during timed calculation (TC) in Exp 1.

| Node Relationships | Mean | SD | P-values | T-values |
|---|---|---|---|---|
| AG-DLPFC | 0.29 | 0.17 | < 0.001 | 5.79 |
| DLPFC-AG | 0.47 | 0.17 | < 0.001 | 7.28 |
| AG-FFG | 0.42 | 0.17 | < 0.001 | 4.72 |
| FFG-AG | 0.22 | 0.17 | < 0.001 | 3.97 |
| AG-IFG | 0.30 | 0.20 | < 0.001 | 6.16 |
| IFG-AG | 0.38 | 0.20 | < 0.001 | 4.25 |
| AG-IPS | 0.26 | 0.16 | < 0.001 | 7.90 |
| IPS-AG | 0.51 | 0.17 | < 0.001 | 6.59 |
| AG-Str | 0.45 | 0.18 | < 0.001 | 5.80 |
| Str-AG | 0.32 | 0.17 | < 0.001 | 6.25 |
| DLPFC-FFG | 0.41 | 0.17 | < 0.001 | 5.95 |
| FFG-DLPFC | 0.26 | 0.17 | < 0.001 | 5.36 |
| DLPFC-IFG | 0.35 | 0.19 | < 0.001 | 6.48 |
| IFG-DLPFC | 0.25 | 0.20 | < 0.001 | 5.44 |
| DLPFC-IPS | 0.52 | 0.16 | < 0.001 | 5.19 |
| IPS-DLPFC | 0.27 | 0.15 | < 0.001 | 3.83 |
| DLPFC-Str | 0.42 | 0.18 | < 0.001 | 5.15 |
| Str-DLPFC | 0.20 | 0.19 | < 0.001 | 4.54 |

| | | | | |
|---|---|---|---|---|
| FFG-IFG | 0.27 | 0.19 | < 0.001 | 2.62 |
| IFG-FFG | 0.41 | 0.21 | < 0.001 | 5.63 |
| FFG-IPS | 0.27 | 0.16 | < 0.001 | 5.21 |
| IPS-FFG | 0.44 | 0.16 | < 0.001 | 6.21 |
| FFG-Str | 0.31 | 0.18 | < 0.001 | 4.18 |
| Str-FFG | 0.26 | 0.19 | < 0.001 | 5.10 |
| IFG-IPS | 0.36 | 0.20 | < 0.001 | 6.16 |
| IPS-IFG | 0.21 | 0.18 | < 0.001 | 8.46 |
| IFG-Str | 0.45 | 0.20 | < 0.001 | 3.98 |
| Str-IFG | 0.34 | 0.19 | 0.013 | 5.58 |
| IPS-Str | 0.36 | 0.17 | < 0.001 | 5.80 |
| Str-IPS | 0.23 | 0.18 | < 0.001 | 6.69 |

Node relationship labels follow "from - to" directionality; all nodes are right hemisphere, comprising angular gyrus (AG), dorsolateral prefrontal cortex (DLPFC), fusiform gyrus (FFG), inferior frontal gyrus (IFG), intraparietal sulcus (IPS), and striatum (Str).

**Supplementary Table 8**
One-sample t-tests of dynamic causal modeling (DCM) and Bayesian model averaging (BMA) node relationship data during timed calculation (TC) in Exp 2 (N=100).

| Node Relationships | Mean | SD | P-values | T-values |
|---|---|---|---|---|
| AG-DLPFC | 0.30 | 0.08 | < 0.001 | > 20 |
| DLPFC-AG | 0.34 | 0.08 | < 0.001 | > 20 |
| AG-FFG | 0.37 | 0.09 | < 0.001 | > 20 |
| FFG-AG | 0.19 | 0.07 | < 0.001 | > 20 |
| AG-IFG | 0.27 | 0.09 | < 0.001 | > 20 |
| IFG-AG | 0.30 | 0.09 | < 0.001 | > 20 |
| AG-IPS | 0.26 | 0.08 | < 0.001 | > 20 |
| IPS-AG | 0.38 | 0.08 | < 0.001 | > 20 |
| AG-Str | 0.38 | 0.09 | < 0.001 | > 20 |
| Str-AG | 0.23 | 0.08 | < 0.001 | 8.60 |
| DLPFC-FFG | 0.41 | 0.08 | < 0.001 | > 20 |
| FFG-DLPFC | 0.24 | 0.08 | < 0.001 | > 20 |
| DLPFC-IFG | 0.36 | 0.08 | < 0.001 | > 20 |
| IFG-DLPFC | 0.19 | 0.08 | < 0.001 | > 20 |
| DLPFC-IPS | 0.29 | 0.08 | < 0.001 | > 20 |
| IPS-DLPFC | 0.36 | 0.08 | < 0.001 | > 20 |
| DLPFC-Str | 0.44 | 0.09 | < 0.001 | > 20 |
| Str-DLPFC | 0.30 | 0.09 | < 0.001 | > 20 |
| FFG-IFG | 0.24 | 0.08 | < 0.001 | > 20 |
| IFG-FFG | 0.33 | 0.09 | < 0.001 | > 20 |
| FFG-IPS | 0.24 | 0.08 | < 0.001 | 9.03 |
| IPS-FFG | 0.34 | 0.07 | < 0.001 | > 20 |

| | | | | |
|---|---|---|---|---|
| FFG-Str | 0.36 | 0.09 | < 0.001 | 8.70 |
| Str-FFG | 0.31 | 0.08 | < 0.001 | > 20 |
| IFG-IPS | 0.23 | 0.09 | < 0.001 | > 20 |
| IPS-IFG | 0.31 | 0.08 | < 0.001 | > 20 |
| IFG-Str | 0.38 | 0.09 | < 0.001 | 9.00 |
| Str-IFG | 0.25 | 0.09 | < 0.001 | 10.00 |
| IPS-Str | 0.41 | 0.09 | < 0.001 | > 20 |
| Str-IPS | 0.20 | 0.07 | < 0.001 | 7.85 |

Node relationship labels follow "from - to" directionality; all nodes are right hemisphere, comprising angular gyrus (AG), dorsolateral prefrontal cortex (DLPFC), fusiform gyrus (FFG), inferior frontal gyrus (IFG), intraparietal sulcus (IPS), and striatum (Str).
